# Spatiotemporal dynamics and modelling support the case for area-wide management of citrus greasy spot in a Brazilian smallholder farming region

**DOI:** 10.1101/839431

**Authors:** F.F. Laranjeira, S.X.B. Silva, R.E. Murray-Watson, A.C.F. Soares, H.P. Santos-Filho, N.J. Cunniffe

## Abstract

Citrus greasy spot (CGS), caused by *Zasmidium citri*, induces premature defoliation and yield loss in *Citrus* spp. CGS epidemiology is well understood in areas of high humidity such as Florida (USA), but remains unaddressed in Brazil, despite differing climatic conditions and disease management practices. We characterize the spatiotemporal dynamics of CGS in the Recôncavo of Bahia, Brazil, focusing on four hierarchical levels (quadrant, plant, grove and region). A survey conducted in 19 municipalities showed that disease is found throughout the entire region with a prevalence (i.e. proportion of affected sampling units) of 100% in groves and plants, and never lower than 70% on leaves. Index of dispersion (*D*) values suggest the spatial pattern of symptomatic units lies somewhere between random and regular. This was confirmed by the parameters of the binary power law for plants and their quadrants (log(*A*)<0 and *b*<1). Variability in disease severity at different plant heights (0.7 m, 1.3 m and 2.0 m) was tested, but no consistent differences were observed. We introduce a simple compartmental model synthesising the epidemiology of the disease, in order to motivate and guide further research. The data we have collected allow such a model to be parameterised, albeit with some ambiguity over the proportion of new infections that result from inoculum produced within the grove *vs.* external sources of infection. By extending our model to include two populations of growers – those who control and those who do not – coupled by the spread of airborne inoculum, we investigate likely performance of the type of cultural controls that would be accessible to citrus growers in Northeastern Brazil. Our model shows that control via removal of the key source of inoculum – i.e. fallen leaves – can be very effective. However, successful control is likely to require area-wide strategies, in which a large proportion of growers actively manage disease.

## Introduction

Citrus greasy spot (*Zasmidium citri* Whiteside), CGS, is considered an important fungal disease in areas of high relative air humidity and temperature (Mondal and Timmer, 2006; Silva *et al.*, 2015; Timmer & Gottwald, 2000; Whiteside, 1988), such as the Caribbean basin and Florida in the USA. The main symptom is irregular leaf spots resembling a brownish-black grease, surrounded by a greenish-yellow halo which is more conspicuous in the disease’s initial stages (Hidalgo *et al.*, 1997). Dark spots – which are leaf symptoms rather than perithecia – are visible on the abaxial side of the leaf and usually correspond to chlorotic areas on the adaxial side. The former increase in size whereas the latter fade away (Timmer & Gottwald, 2000; Whiteside, 1988). *Z. citri* also infects fruits causing unsightly rind blemishing which can affect trade value (Timmer & Gottwald, 2000). Perithecia are produced only in fallen decomposing leaves (Timmer *et al.*, 2000). CGS leads to premature defoliation followed by reduction of both fruit size and plant vigour (Timmer *et al.*, 2000). At least in Cuba, associated loss of yield can reach 5 t/ha (Diaz *et al*, 1985).

Despite its importance in many citrus producing regions (Timmer *et al.*, 2000), CGS is considered to be only a minor disease in Brazil. Few studies characterizing its epidemiology in the Brazilian context have therefore been performed (Laranjeira *et al.*, 2005). However, CGS is important in Bahia State, the second most important citrus growing region in Brazil. A regional survey in Bahia showed that the disease occurs at high prevalence (100% of the sampled groves) meaning CGS is endemic, in the sense of being regularly found and very common in the area (Silva *et al.*, 2009). Moreover, CGS has been shown to exert a severe defoliating effect on cultivated citrus in Bahia, with one study estimating that 500 leaves are lost due to the disease per year from each infected “Pera” sweet orange plant (Silva *et al.*, 2015). Surprisingly, however, our discussions with citrus growers in Bahia reveal they do not perceive CGS to be a yield-limiting factor, and recent reports indicate that less than 4% of growers try to manage the disease (Rodrigues, 2018).

Studies based on the spatiotemporal patterns of the disease may help to better understand the pathogen’s dispersion mechanism(s) and to develop additional strategies for disease management. In particular, understanding spatial patterns of disease offers the promise of understanding the balance between auto- and allo-infection at the grove scale, and in turn of revealing the extent to which disease control must be attempted on an area-wide basis. The association of the life-cycle with weather is well studied (Garcia *et al.*, 1980; Hidalgo *et al.*, 1997; Mondal & Timmer, 2003a) but there are no reports regarding spatial patterns.

In Cuba, rain and relative air humidity are correlated with CGS incidence. The disease builds up between summer and early autumn, when the most intense rains occur and the relative humidity increases to between 84% and 90% (Garcia *et al.*, 1980). In Costa Rica ascospore release closely follows seasonal rain patterns. The number of trapped ascospores quickly rises in May, peaking at the beginning of June (late spring). This is then followed by considerable decline in July and the number of released ascospores remains low for the rest of the year (Hidalgo *et al.*, 1997). In Florida in the USA the pseudothecia grow slowly on the decomposing litter during the relatively dry spring, and ascospore release is retarded until the summer rains, with symptoms following only in the winter (Timmer *et al.*, 2000). In Northeastern Brazil (Bahia), however, the weather conditions are conducive for inoculum production in all seasons. The relative humidity is always higher than 70% and rain events occur throughout the year (Silva *et al.*, 2015).

Despite the amount of information generated, no study has attempted to construct a mathematical model of CGS. In this paper, we develop a model of *Z. citri* population dynamics which we use to compare the likely efficacy of disease management strategies. In Florida and other regions with an extensive citrus-production industry, cultivation tends to be based around large commercial operations, within which *Z. citri* is controlled by fungicides (Mondal & Timmer, 2006a). However, small-holder growers are in the majority in Bahia, and such growers do not have ready access to expensive agrochemicals, or even to the machinery required to apply such products. We therefore concentrate here on the performance of a cultural control of the type that could potentially be performed by resource-poor growers. We focus in particular on removal of fallen leaf litter, a management strategy originally proposed by Whiteside (1970) nearly 50 years ago. Such a localised cultural control done by an individual grower can clearly only affect the component of the epidemic spread driven by within-grove production of inoculum. We therefore particularly focus on using our model to assess whether and under which conditions the performance of cultural disease management can be improved when it is taken up area-wide by a significant fraction of a community of growers (Bassanezi *et al.*, 2013; Bergamin Filho *et al.*, 2016; Sherman *et al.*, 2019).

## Materials and Methods

### Levels of the spatial hierarchy and sampling procedures

We divided our sample space into four levels: plant quadrants (Hierarchical Level 1, HL1), plants in a grove (HL2), groves (HL3) and the region (HL4). A plant quadrant was considered as each side of a plant, i.e. there were two within-row quadrants and two across-rows quadrants. A grove was defined as a uniform production unit inside a farm; groves we considered ranged in size from 1 Ha to 3 Ha. Plants in a single grove invariably had the same rootstock and scion varieties, as well as the same age (which varied between three and six years over the set of groves we considered) and planting practice (i.e. spacing within and across rows).

To guide decisions about logistics and methods used at the other spatial levels, CGS prevalence (fraction of disease cases in a sample of a given region) and incidence (fraction of affected plants in a given orchard) were first quantified in our HL4, the ‘Recôncavo da Bahia’ region (an area formed of 20 munipalities in the vicinity of Todos os Santos Bay on the East coast of Brazil near the city of Salvador; see also Fig. 1). As a first criterion, only the 19 municipalities with at least 100 ha of citrus groves were sampled (IBGE, 2017): Cachoeira (38° 56’ 56”W, 12° 35’ 26”S), Cabaceiras do Paraguaçu (39° 11’ 12”W, 12° 37’ 50”S), Castro Alves (39° 14’ 45”W, 12° 41’ 32”S), Conceição da Feira (38° 58’ 31”W, 12° 30’ 49”S), Conceição do Almeida (39° 10’ 50”W, 12° 48’ 03”S), Cruz das Almas (39° 08’ 35”W, 12° 40’ 59”S), Dom Macedo Costa (39° 12’ 23”W, 12° 53’ 06”S), Governador Mangabeira (39° 02’ 33”W, 12° 35’ 34”S), Jaguaripe (39° 10’ 14”W, 13° 11’ 41”S), Maragogipe (39° 02’ 29”W, 12° 45’ 04”S), Muniz Ferreira (39° 06’ 44”W, 12° 59’ 50”S), Muritiba (39° 09’ 44”W, 12° 37’ 09”S), Sapeaçu (39° 11’ 32”W, 12° 38’ 28”S), São Félix (39° 02’ 33”W, 12° 42’ 22”S), Santo Amaro da Purificação (38° 50’ 54”W, 12° 32’ 27”S), São Felipe (39° 02’ 50”W, 12° 44’ 00”S), Santo Antônio de Jesus (39° 18’ 51”W, 12° 58’ 34”S), São Miguel das Matas (39° 27’ 58”W, 12° 58’ 45”S) and Varzedo (39° 20’ 44”W, 12° 58’ 40”S). One grove per municipality was sampled; each was no larger than 3 Ha, and consisted of ‘Pera’ sweet orange grafted on Rangpur lime. The choices of evaluation method, number of groves per municipality, citrus variety and grove size were based on CGS sampling procedures previously recommended for the Recôncavo Baiano region (Silva *et al.*, 2009). In each grove a ‘W’-shaped path was followed in which 30 plants were arbitrarily chosen and thoroughly visually evaluated for the presence of typical CGS symptoms. CGS prevalence in the region was then calculated as the proportion of groves with affected plants, whereas incidence was estimated as the proportion of symptomatic plants in each grove.

**Figure 1.**
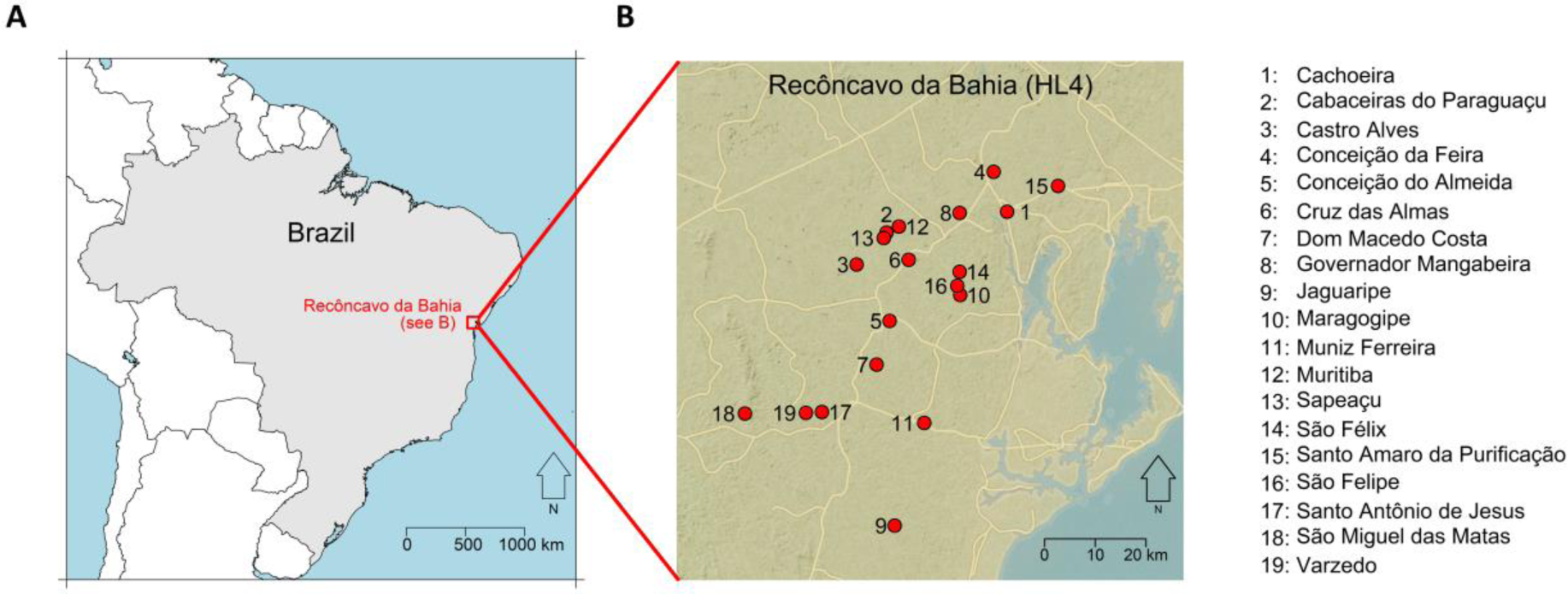
Locations in Recôncavo of Bahia (our Hierarchical Level 4) that were sampled for disease. Only the 19 (of the 20) municipalities in the region with at least 100 ha of citrus groves were sampled. (A) Location of the region in Brazil. (B) The 19 locations which were sampled. The maps were produced using the R packages maps (Becker et al., 2018) and OpenStreetMap (Fellows, 2019).

Data thus collected were the basis for examining the spatiotemporal patterns in HL1, HL2 and HL3. Ten groves of ‘Pêra’ sweet orange grafted on Rangpur lime (ranging from six to ten years old) were selected in three separate subregions of the Cruz das Almas municipality. Following a ‘W’-shaped path, 30 plants were inspected in each grove. The number of inspected plants in each leg of the path was chosen according to the grove’s shape and size. The position of each plant in each leg of the path was chosen before starting the procedure in each grove. The first five mature leaves from five different branches were examined in each plant’s quadrant (two within planting rows and two between rows) in search of typical symptoms. The proportions of symptomatic leaves, quadrants and leaves were then calculated. The evaluations began in August 2006 and were perfomed monthly until February 2008.

### Temporal patterns

Monthly mean incidence (proportion of symptomatic plants, quadrants or leaves) was calculated by averaging data from the ten sampled groves and used to plot disease progress curves.

### Weather data and distributed lag analysis

The following variables were recorded daily by the Embrapa Cassava & Tropical Fruits weather station, 10 km east of the three evaluated groves: average rain (mm); days of rain each month; minimum, mean and maximum air temperature (°C) and mean air relative humidity (%). Data were organized in a monthly set and a distributed lag analysis was performed against *psl* (i.e. the proportion of symptomatic leaves) considering time lags of up 6 months. Distributed lag analysis is a regression used to predict current values of a dependent variable based on both the current value of an explanatory variable and the lagged (past period) values of that same variable (Laranjeira *et al.*, 2003; Chatfield, 2004; Paul *et al.*, 2007).

### Spatial pattern and dynamics

To examine relations between the distinct levels of our spatial hierarchy, HL1 (among quadrants of a plant), HL2 (plants of a grove) and HL3 (groves in the region), the index of dispersion (*D*) and the Binary Power Law were used (Madden & Hughes, 1995). The sampling units (*N*) were respectively the plant quadrants (HL1), the plants (HL2) and the groves (HL3); and the potential diseased entities (*n*) were leaves (HL1 and HL2) and plants (HL3). The hierarchical levels had the following combinations of *N, n*: 1200, 5 (HL1); 300, 20 (HL2); 10, 30 (HL3).

The observed variance (V_obs_) as well as the expected binomial variance (V_bin_) were calculated for each hierarchical level and evaluation date (Madden & Hughes, 1995; Madden, Hughes & Vandenbosch, 2007), in order to obtain the index of dispersion (*D* = V_obs_ / V_bin_). A χ^2^ test was used to assess whether there was any departure from randomness. The index of dispersion is used to assess the spatial pattern of a single hierarchical level on a given evaluation date. Values of *D* higher than 1 were considered indicators of aggregation of symptomatic plants, those lower than 1 were taken to reflect a regular pattern. Those values which did not statistically differ from 1 were an indication of randomness, that is, a distribution of symptomatic plants without any regularity or aggregation.

The Binary Power Law, an adapted form of the Taylor Power Law for proportional data where variances do not increase monotonically with means (Madden *et al.*, 2018), was used to assess the spatial heterogeneity of disease incidence. If the disease incidence was found to be aggregated (indicated by a slope, *b*, < 1), this would favour the hypothesis that auto-infection was an important driver of CGS incidence in groves. However, if the disease incidence was uniform (*b* > 1), this would suggest that allo-infection contributed more to CGS incidence.

Whilst the index of dispersion considers individual datasets, the Binary Power Law (BPL) can be used to assess multiple data sets (Madden *et al.*, 2018). The BPL uses observed variance and expected binomial variance to estimate the spatial heterogeneity of CGS.

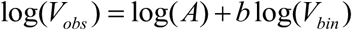

A suitable *F* test was used to determine the significance of the relations between log(V_bin_) and log(V_obs_) (Statistica 5.0, Tulsa, USA); goodness-of-fit was assessed via *R*^2^ and an analysis of residual patterns *vs.* the expected values of log (V_bin_) (Madden & Hughes, 1995); Residual normality was also tested (Looney & Gulledge, 1985). The parameters corresponding to the intercept (log*(A)*) and the regression slope (*b*) were considered significant if different from 0 and 1, respectively (t test, *p* < 0.05) (Madden & Hughes, 1995). The binary power law is used to assess the spatial pattern of a collection of evaluation dates in a given hierarchical level. A value *b* > 1 was taken as an indication of underdispersion, whereas *b* < 1 indicated overdispersion and *b* = 1 randomness.

### Vertical pattern

The variability in the proportion of symptomatic leaves (*psl*) among plant heights was evaluated in three randomly selected groves in July 2007 (these three groves were not part of the set of groves used to assess spatial patterns). Leaves at three heights, 0.7 m; 1.3 m and 2.0 m (treatments) were sampled in each of 30 plants (replications) per grove (block) in a total of 270 sampling units. For each of these units, 20 leaves were evaluated and the number of infected leaves was scored. These data were analysed using a generalized linear mixed model for binomial counts via the R package `lme4` (Bates *et al.*, 2015), taking the grove as a (fixed) blocking factor, height as a (fixed) treatment effect and plant as a random effect. Significance was assessed by comparing nested models via Likelihood Ratio tests (Lewis, Butler & Gilbert, 2011).

### Mathematical model

We develop a simple compartmental model of the epidemiology of *Z. citri*, focusing on only the key features relevant to CGS epidemiology in Brazil (Fig. 2). Our core “single-grower” model tracks the disease status of individual leaves on a representative plant within a single grove, since this is the scale at which disease symptoms are expressed and at which symptoms were scored in the survey. However, since the disease status of individual plants sets the grove-scale dynamics, our model produces results of relevance to our Hierarchical Level 3 (i.e. an individual grove). The single-grower model is then extended to allow for area-wide control (see Area-wide disease management, below), by accounting for disease spread within two coupled populations of growers, those who control disease and those who do not.

**Figure 2.**
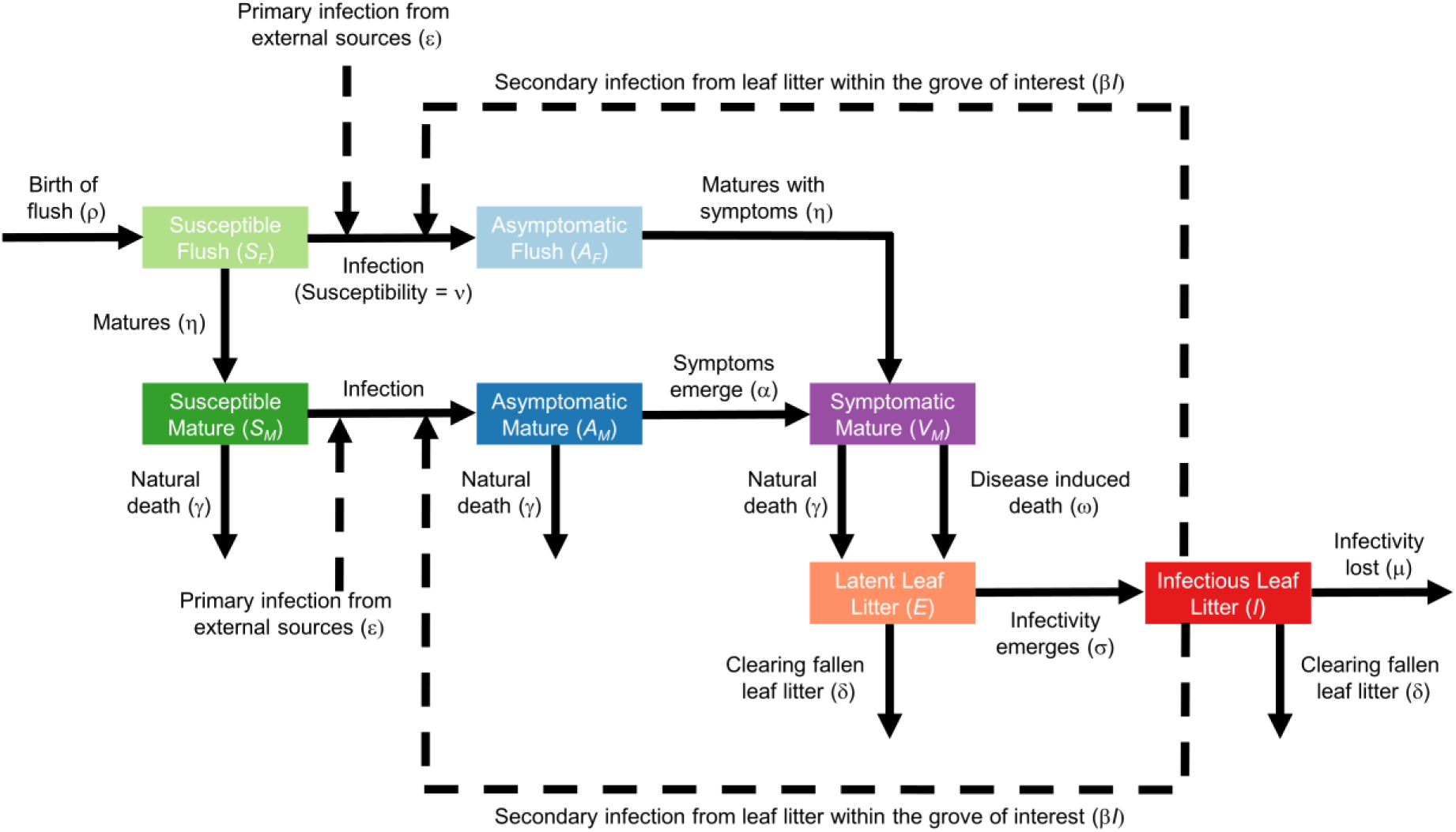
Schematic showing the structure of the single-grower mathematical model for within- and between-grove spread of citrus greasy spot.

We separate newly-created flush leaves from mature leaves in our model, since we scored only mature leaves for disease symptoms. We assume that flush transitions to maturity after an average of 1/η units of time, where η is the rate of maturation. We also assume that infected leaves do not show symptoms immediately, with the incubation period of mature leaves assumed to average 1/α units of time, where α is the rate of development of symptoms. We furthermore assume that only mature leaves can exhibit symptoms, with infected flush leaves starting to show symptoms immediately after reaching maturity (Fig. 2, Table 1).

**Table 1.** Meaning of symbols and parameters, as well as default parameter values, in the mathematical model of disease experienced by a single-grower (Fig. 1). Sources are given for model parameters (see also Results).

| Symbol | Meaning | Value | Source |
| --- | --- | --- | --- |
| $S_F$ | Number of susceptible (healthy) flush leaves | Varies | n/a |
| $A_F$ | Number of infected (asymptomatic) flush leaves | Varies | n/a |
| $S_M$ | Number of susceptible (healthy) mature leaves | Varies | n/a |
| $A_M$ | Number of infected (asymptomatic) mature leaves | Varies | n/a |
| $V_M$ | Number of infected (symptomatic) flush leaves | Varies | n/a |
| $E$ | Number of infected fallen leaves that are not yet sporulating | Varies | n/a |
| $I$ | Number of infected fallen leaves that have started sporulating | Varies | n/a |
| $t$ | Time (in months) | Varies | n/a |
| $\rho$ | Birth rate of flush | 6000 month<sup>-1</sup> | Turrell (1961) |
| $\beta$ | Rate of secondary infection from infected fallen leaves | Fitted to data | See Fig. 9 |
| $\varepsilon$ | Rate of primary infection from outside focal grove | Fitted to data | See Fig. 9 |
| $\nu$ | Proportionate susceptibility of flush vs. mature | 1 | Mondal & Timmer (2003b) |
| $\eta$ | Rate of maturation of flush to become mature | 1/3 month<sup>-1</sup> | Spiegel-Roy & Goldschmidt (1996) |
| $\gamma$ | Rate of death of mature leaves | 1/9 month<sup>-1</sup> | Wallace A, Zidan ZI, Mueller RT (1954)<br>Spiegel-Roy & Goldschmidt, (1996) |
| $\alpha$ | Rate of emergence of symptoms | 1/2 month<sup>-1</sup> | Mondal & Timmer (2006b) |
| $\omega$ | Rate of disease-induced death | 2/9 month<sup>-1</sup> | Timmer *et al.* (2000) |
| $\sigma$ | Rate of emergence of infectivity on infected fallen leaves | 1/2 month<sup>-1</sup> | Mondal & Timmer (2006b) |
| $\mu$ | Rate of loss of infectivity on fallen leaves | 1 month<sup>-1</sup> | Mondal & Timmer (2003b) |
| $\delta$ | Rate at which leaf litter is cleared | 0 month<sup>-1</sup><br>(scanned over in assessing controls) | Litter clearance is not common practice in Brazil |

A total of five leaf classes are therefore tracked: susceptible (i.e. healthy) flush (*S*_*F*_), asymptomatic infected flush (*A*_*F*_), susceptible mature leaves (*S*_*M*_), asymptomatic infected mature leaves (*A*_*M*_) and symptomatic mature leaves (*V*_*M*_). Since only mature leaves were scored for disease, the proportion of symptomatic leaves (*psl*) as recorded in the survey data corresponds to Γ = *V*_*M*_/(*S*_*M*_*+A*_*M*_*+V*_*M*_). Mature leaves are assumed to abscise and fall to the ground at rate γ; symptomatic infected leaves are assumed to suffer additional disease-induced mortality at rate ω (Mondal & Timmer, 2006a). This continual turnover of infected and uninfected leaves is offset by production of new flush; for simplicity we assume production occurs at constant rate ρ (Laranjeira *et al.*, 2003)

Secondary infection within the grove of interest occurs at rate β via sporulation off of fallen leaf litter that was once an infected mature leaf (Mondal & Timmer, 2006b; Mondal *et al*., 2003; Timmer *et al*., 2000); this follows a latent period on the ground which averages 1/σ units of time. Tracking this within-grove secondary infection pathway requires us to track a further two classes of leaf litter: latently infected (*E*) and sporulating infectious (*I*). The only source of latently infected leaves is mature symptomatic (i.e. leaves in class *V*_*M*_) abscising from the tree; i.e. any pre-symptomatic infected leaves that abscise are assumed to not lead to infectious litter, and there is no spread of the pathogen within the litter (Fig. 2, Table 1).

We assume that sporulating litter remains infectious for an average of 1/μ units of time, and that litter is potentially cleared by the grower at rate δ. We also allow for primary infection from sources of inoculum outside the grove of interest. In our single-grower model we assume this occurs at a constant rate (ε). We allow the susceptibility of flush to differ from that of mature leaves: the proportionate susceptibility of flush is assumed to be ν (with an equal effect on susceptibility to infection via both the primary and the secondary infection pathways) (Fig. 2, Table 1).

The system of equations defining the single-grower model is therefore as follows.

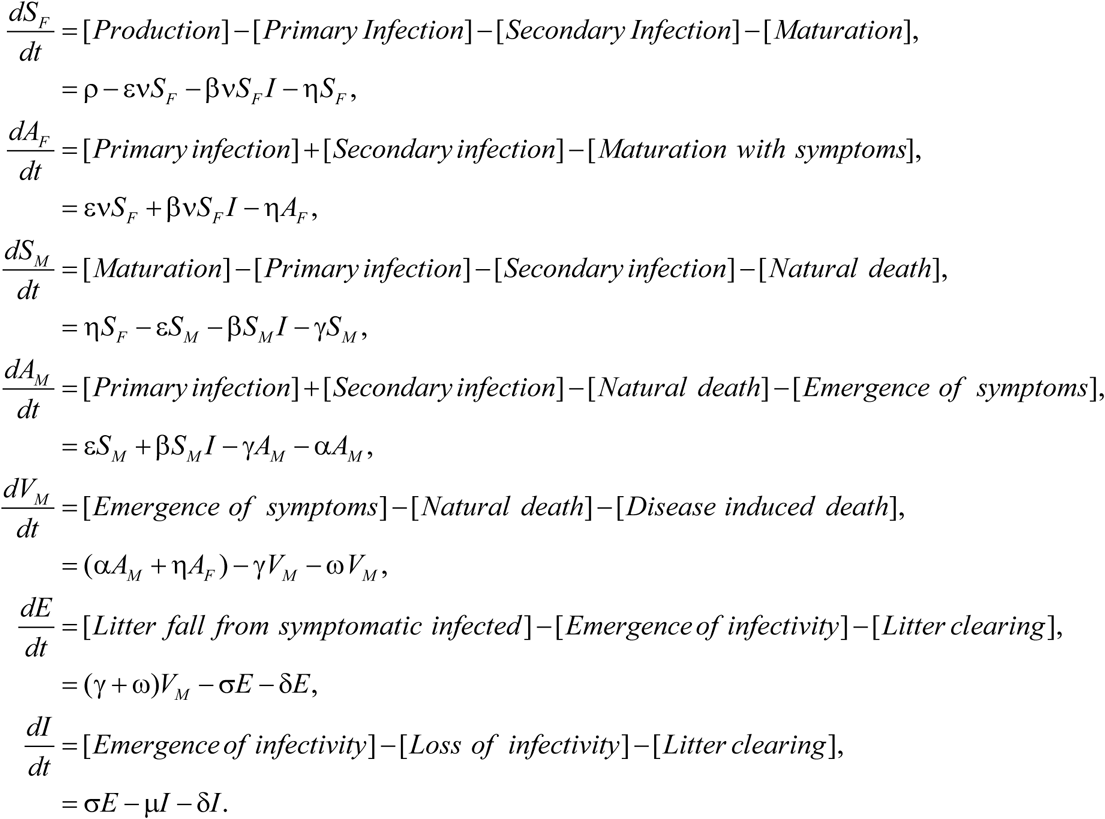

A reference implementation of the model in the programming language R (R Core Team, 2018) is available at https://github.com/nikcunniffe/Citrus-Greasy-Spot; it relies on the R package `deSolve` (Soetaert *et al.*, 2010) for numerical solution of ordinary differential equations.

### Area-wide disease management

A key simplification in our model is that the rate of primary infection is held constant. For disease management attempted by a single grower in isolation this is a reasonable assumption, inasmuch as the primary infection pathway is totally unaffected by control done within the grove. However, if other growers were also to attempt to control disease, this would clearly lead to a reduction in the amount of exported inoculum by these growers. In turn this would lead to a reduction in rates of primary infection across the entire region, since all growers would then suffer less infection. To account for this feedback between the local intensity of disease control and area-wide rates of primary infection, we extended the initial mathematical model to include two populations of grower: controllers (C) and non-controllers (NC). This modified model allows us to understand the performance of area-wide management.

We assume these two groups differ in how they manage CGS; the controllers undertake localised control and clear leaf litter at rate δ. The non-controllers do not clear litter (i.e. have δ fixed at zero). We also assume that there is no spatial structure, thus all growers in the region of interest within each class are homogenous in their control practices and grove size. The model therefore tracks the average disease dynamics experienced by a typical grower in each class. For simplicity we furthermore assume that all inoculum causing primary infection is generated from somewhere within the modelled region, i.e. there is no very long-distance spread from outwith the set of groves tracked by our model or from more local sources of inoculum within the region such as non-cultivated citrus.

Under this scenario, the rate of primary infection (ε) is no longer constant but should vary as ε = ς (*πI*^*C*^ + (1-*π*) *I*^*NC*^) (Fig. 3), in which π is the proportion of growers controlling disease and *I*^*C*^ and *I*^*NC*^ correspond to the levels of infection in typical groves of controllers and non-controllers, respectively. To ensure the results of the area-wide model are consistent with the underlying single-grower model, we set the constant of proportionality, ς, as ς = ε / *I*_*∞*_, where *I*_*∞*_ is the equilibrium level of infection in the absence of control and ε is the rate of primary infection, both lifted from the core single-grower model.

**Figure 3.**
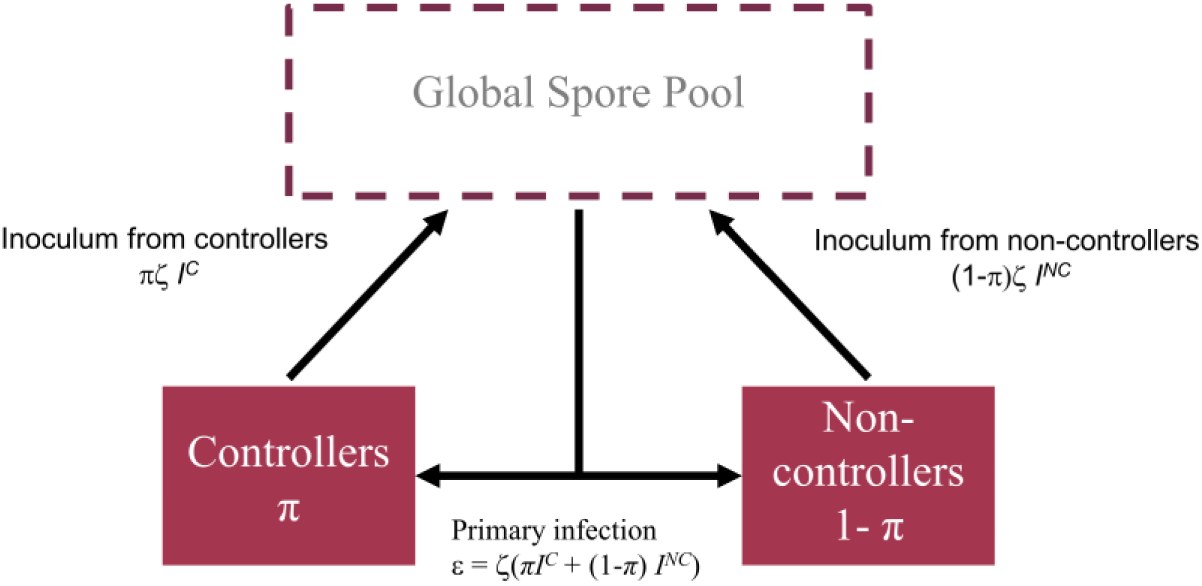
Schematic showing the structure of the mathematical model for area-wide implementation of control. The contribution of a grove to the area-wide primary infection rate depends on its control status. Inoculum from both types of growers mixes in the global spore pool and falls on any single grove of either type in equal measure.

The new model for the controllers is defined as follows, with terms that differ from the single-grower model – or which differ between controllers and non-controllers – highlighted in red. Note the additional superscripts on the state variables, e.g. 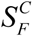 corresponds to *S*_*F*_ for growers which control disease (the corresponding state variable for the non-controllers is 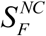).

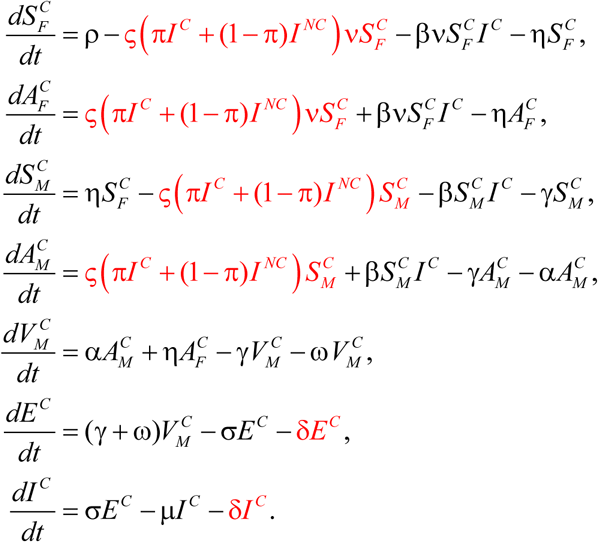

The equations for growers who do not control disease are very similar:

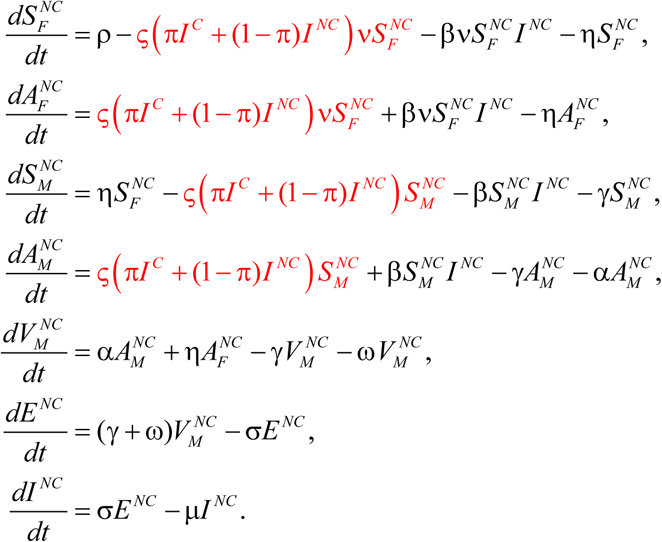

The only qualitative difference between the models for the two types of grower is that the δ*I* and δ*E* terms are absent from the final equation for the non-controlling growers.

## Results

### Spatial variability among plant heights

The proportion of symptomatic leaves at individual heights (0.7m, 1.3m, 2.0m) on individual plants varied between 0.70 (i.e. 14 out of 20) and 1.0 (i.e. 20 out of 20), with overall mean 0.90. There was a significant interation between grove and height (Likelihood Ratio Test: 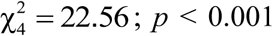), with an increase in disease severity with height in the most heavily infected grove, but no increase in the other two groves. Since there was only a response to height in one out of the three groves tested, we conclude that there was no reliable effect of height on disease severity (see also Supplementary Text 1).

### Prevalence and Incidence in Recôncavo da Bahia

Typical symptoms of CGS were found in all sampled groves, leading to a 100% prevalence. Symptoms were also detected on every assessed plant in each single grove, leading to a 100% incidence.

### CGS temporal pattern

CGS symptoms were observed in 100% of evaluations, groves, plants and plant quadrants in each of the ten monthly sampled areas in the Cruz das Almas municipality. The only variability in these results was detected for the proportion of symptomatic leaves (*psl*), which increased over 19 months, ranging from 0.77 in August 2006 to a maximum of 0.96 in October 2007 (Fig. 4). Although there was some systematic linear increase in the incidence, oscillations were found. However, there were insufficient data to test for periodic behaviour in *psl*, e.g. via spectral analysis.

**Figure 4.**
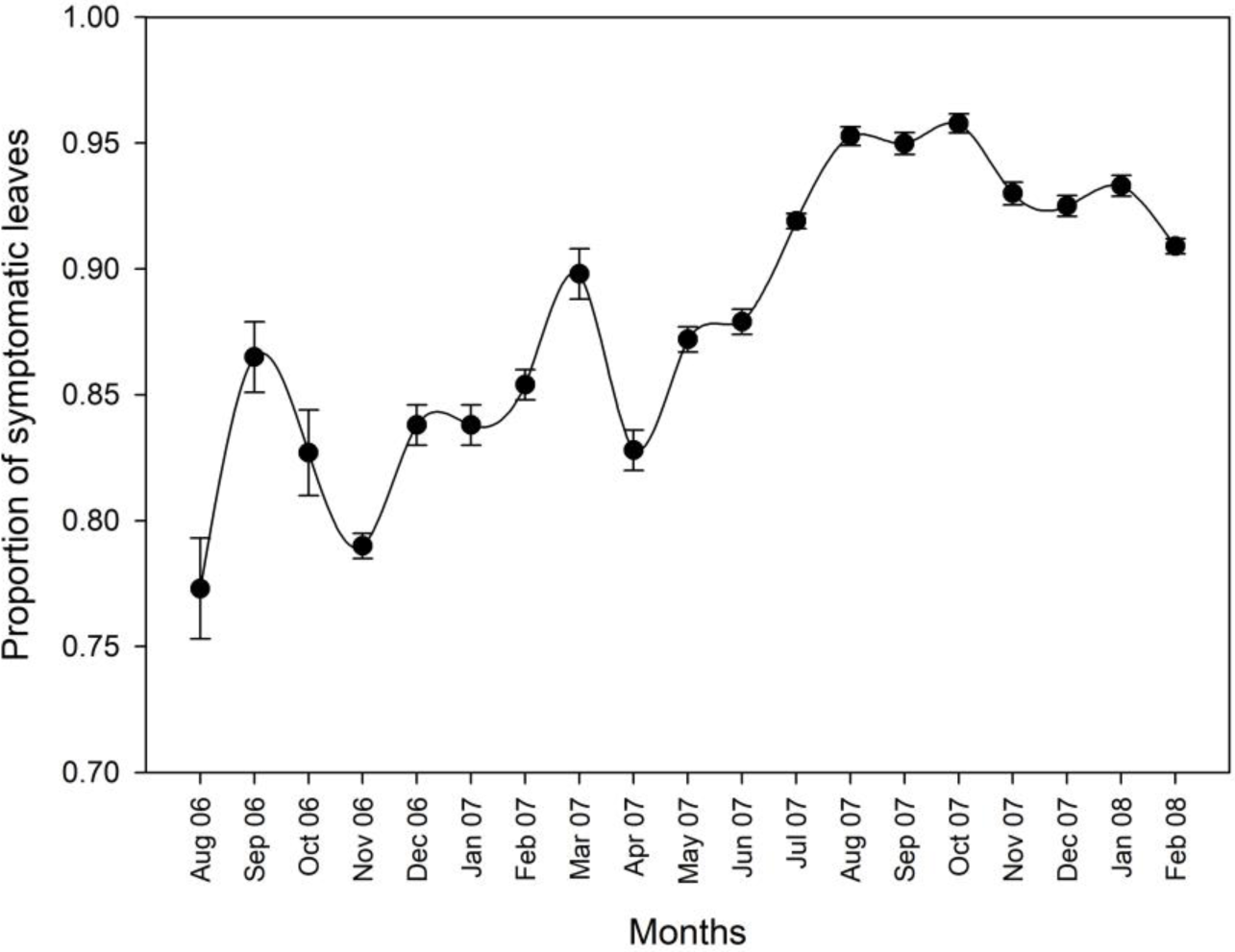
Progress of citrus greasy spot measured as proportion of symptomatic leaves in ten sweet orange groves between August 2006 and February 2008 in Cruz das Almas, Recôncavo Baiano, Brazil. Bars represent standard error.

### Weather variables

Distributed lag analysis performed between the weather variables and CGS *psl* did not reveal any significant relationship (Fig. 5).

**Figure 5.**
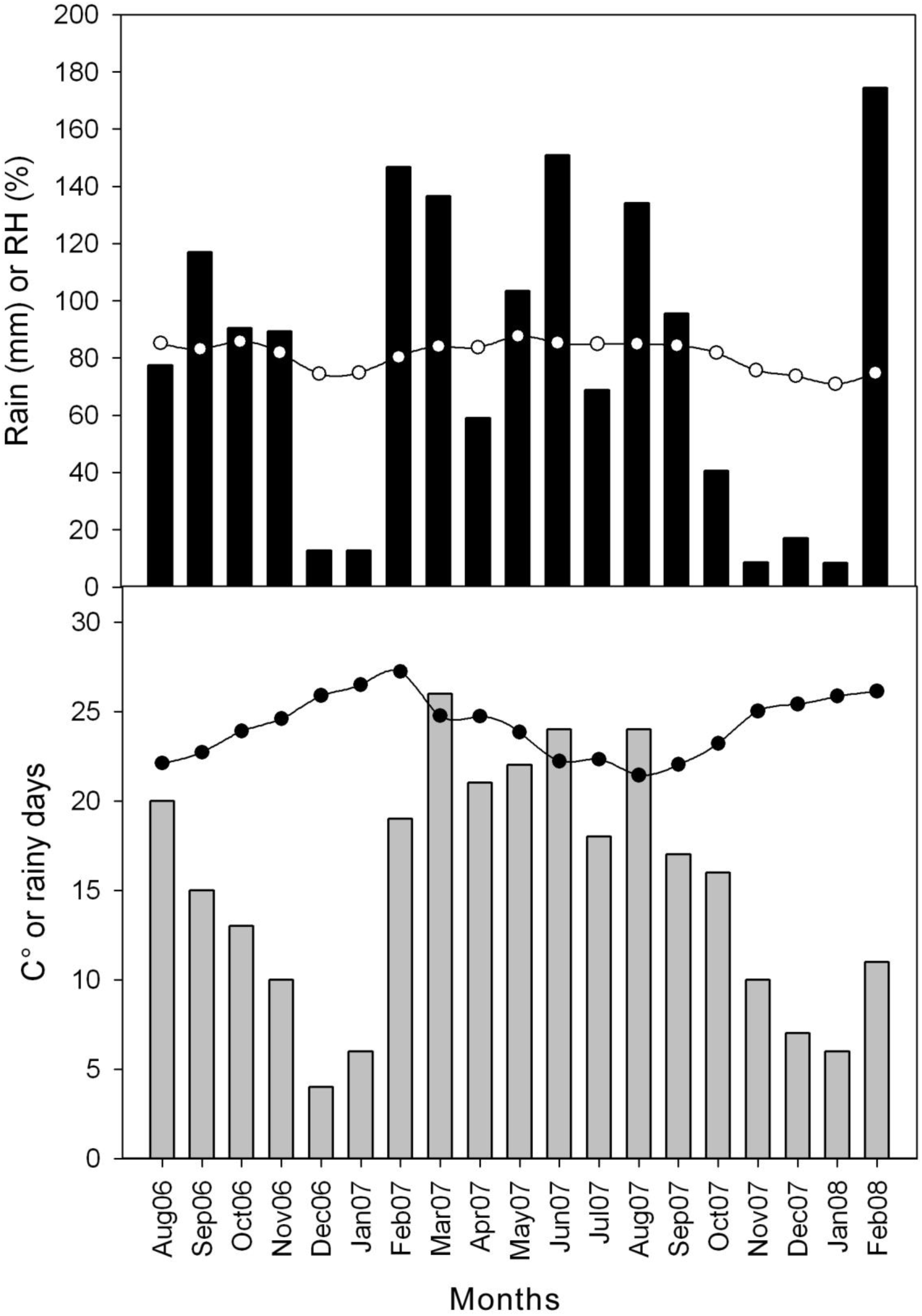
Mean monthly values for weather variables in Recôncavo Baiano between August 2006 and February 2008. Lines are relative air humidity or temperature, bars area rain or number of rainy days.

### Spatial pattern and dynamics

It was not possible to perform any analysis of data from HL3 (among groves) and HL4 (among municipalities) due to lack of variability. In HL1 (plant quadrants) 58% of *D* values were statistically similar to 1 (randomness), whereas 2% were above 1 (aggregation), and 40% below 1 (regularity) (Fig. 6A). At HL2 (plants), the index of dispersion (*D*) had 76%, 12% and 12% of observations statistically equal to, higher than and lower than 1, respectively (Fig. 6B). No relationship between *D* and the observed range of CGS *psl* could be observed, for plant quadrants or plants (Fig. 6). The predominance of *D* values ≤ 1 for HL1 and HL2 could be confirmed by temporal dynamics of *D* means (Fig. 7) and the very low standard errors reinforce this point.

**Figure 6.**
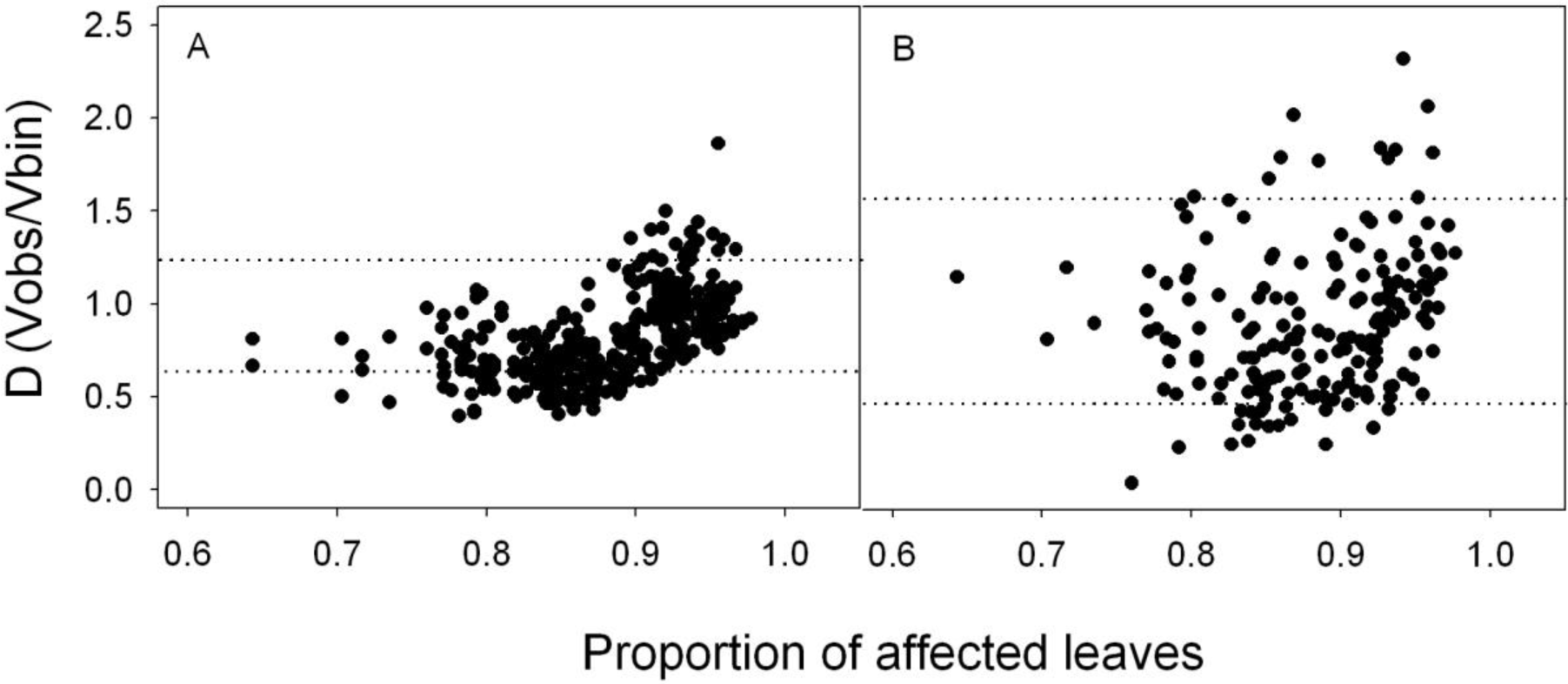
Relationship between citrus greasy spot incidence in leaves and binomial index of dispersion (D) for quadrants (A) and plants (B), in ten sweet orange groves of Recôncavo Baiano, Brazil. Horizontal dotted lines indicate the randomness upper and lower limits for each spatial level. Ds indicate aggregation when above the upper limit and regularity when below the lower limit.

**Figure 7.**
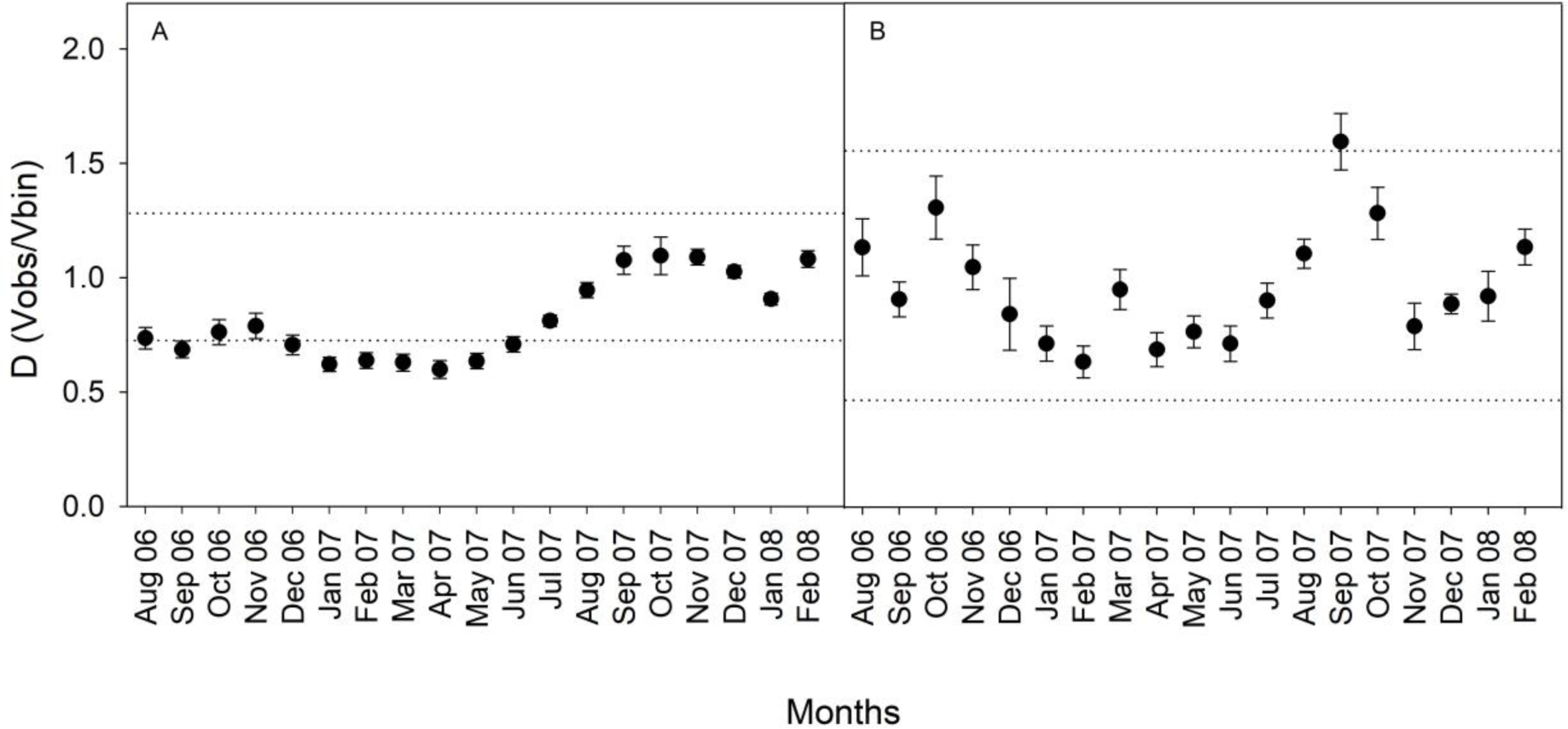
Binomial index of dispersion (D) dynamics for citrus greasy spot in quadrants (A) and plants (B) in ten sweet orange groves in Recôncavo Baiano, Brazil. Values not different from 1 indicate randomness; values above 1 indicate aggregation and those below 1, regularity. Vertical bars indicate standard error.

The binary power law was also used to analyse the CGS spatial patterns in HL1 and HL2. The regression between log (V_bin_) and log (V_obs_) was highly significant for both spatial levels (Fig. 8). The residuals were randomly distributed, but in both cases the coefficients of determination were below 0.7. The regression parameters, log(*A*) and *b*, were significantly lower than 0 and 1, respectively [(log(A): *t* (380)= −16.2, *p* < .00001)), (b: *t* (380)= 28.7, *p* < .0001)] for HL1 (quadrants) and HL2 (plants) [(log(A): *t* (190)= −4.4, *p* < .00001)), (b: *t* (190)= 13.1, *p* < .00001)] (Fig. 8).

**Figure 8.**
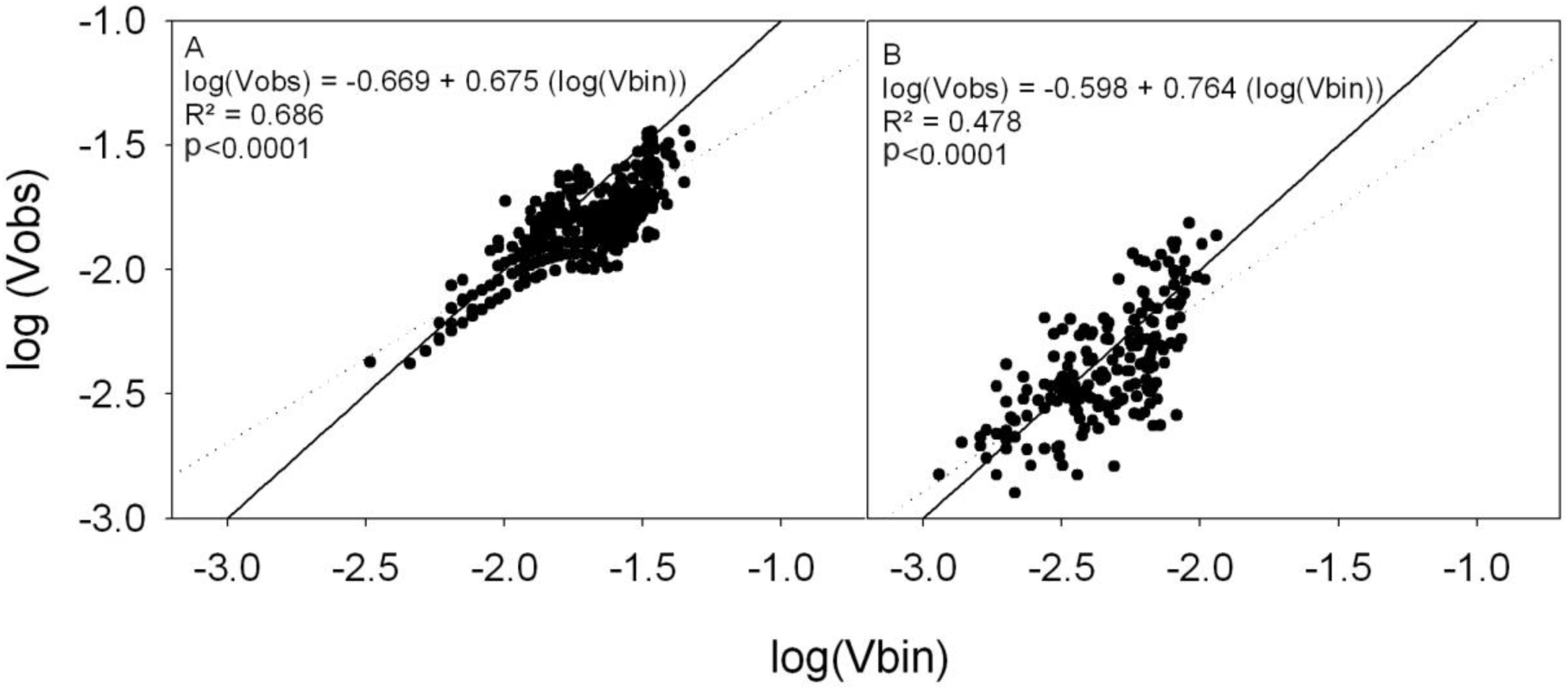
Binary power law. Relationship between the logarithms of observed variance and binomial variance for citrus greasy spot in quadrants (A) and plants (B) in ten sweet orange groves in Recôncavo Baiano, Brazil. Continuous line represents the expected relationship under randomness and the dotted line, the actual regressions. The regressions were significant (p<0.0001) and the parameters log(A) and b were significantly lower than 0 and 1, respectively for both cases.

### Model parameterisation

We assume that leaves mature after approximately three months (Spiegel-Roy & Goldschmidt, 1996), and so set the rate at which flush transitions to become mature leaves to be η = 1/3 month^-1^. We furthermore assume that the average lifetime of a leaf is approximately one year (Wallace *et al*. 1954, Spiegel-Roy & Goldschmidt, 1996), which – after accounting for an average of three months spent as flush – means that the rate of natural death of mature leaves should be γ = 1/9 month^-1^. Figure 1 of Turrell (1961) indicates that a ten-year old citrus tree would have approximately 70,000 leaves. Taking this as a rough estimate of the number of leaves on a mature tree in the absence of disease – predicted to be 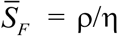 and 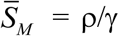 as the steady state of the system of differential equations for the single-grower model when there is no disease − allows us to fix the rate of production of flush to be ρ = 6,000 month^-1^ (where, in the light of the approximate nature of our calculation, we have rounded this estimate to only a single significant figure).

The incubation period between infection of a mature leaf and first emergence of symptoms is approximately two months (Mondal & Timmer, 2006a), and so we take the rate of emergence of symptoms on mature leaves to be α = 1/2 month^-1^. Since the average lifetime of a diseased leaf is approximately three months (Timmer & Gottwald, 2000; Timmer *et al.*, 2000), 1/(γ + ω) should be 3 months, which given our earlier estimate for the value of γ allows us to set the rate of disease-induced death to be ω = 2/9 month^-1^. Data presented in Mondal & Timmer (2002) suggest it takes approximately two months for sporulation to start after an infected leaf falls to the ground, and that – under constant conditions – sporulation would cease after three months. We therefore take the rate at which pre-infectious leaf litter enters the infected class to be σ = 1/2 month^-1^, and the rate of loss of infectivity of infectious leaf litter to be μ = 1 month^-1^ (since litter is actively producing spores for an average of one month).

We assume that recently emerged and mature leaves are equally susceptible (Mondal & Timmer, 2006a), and so take the proportionate susceptibility to be ν = 1. Noting that clearing fallen litter is uncommon in Bahia, we take the default rate of clearance to be δ = 0 (although we scan over values of this parameter when we model disease management; see below).

### Scenarios for primary and secondary infection

The only remaining model parameters that remain to be fixed are the rates of primary (ε) and secondary (β) infection. We identify reasonable values of these parameters by searching for pairs of values leading to *psl* of Γ = 0.85 when the model has reached its disease-present equilibrium, to match the data taken in our survey (*cf.* Fig. 4). Given we are estimating a pair of parameters (ε and β) from a single datum (equilibrium *psl*), there are an infinite number of pairs of parameter values which lead to the desired equilibirium severity (marked by the black curve on Fig. 9A). Our current data do not allow us to distinguish the values of these parameters in more detail.

**Figure 9.**
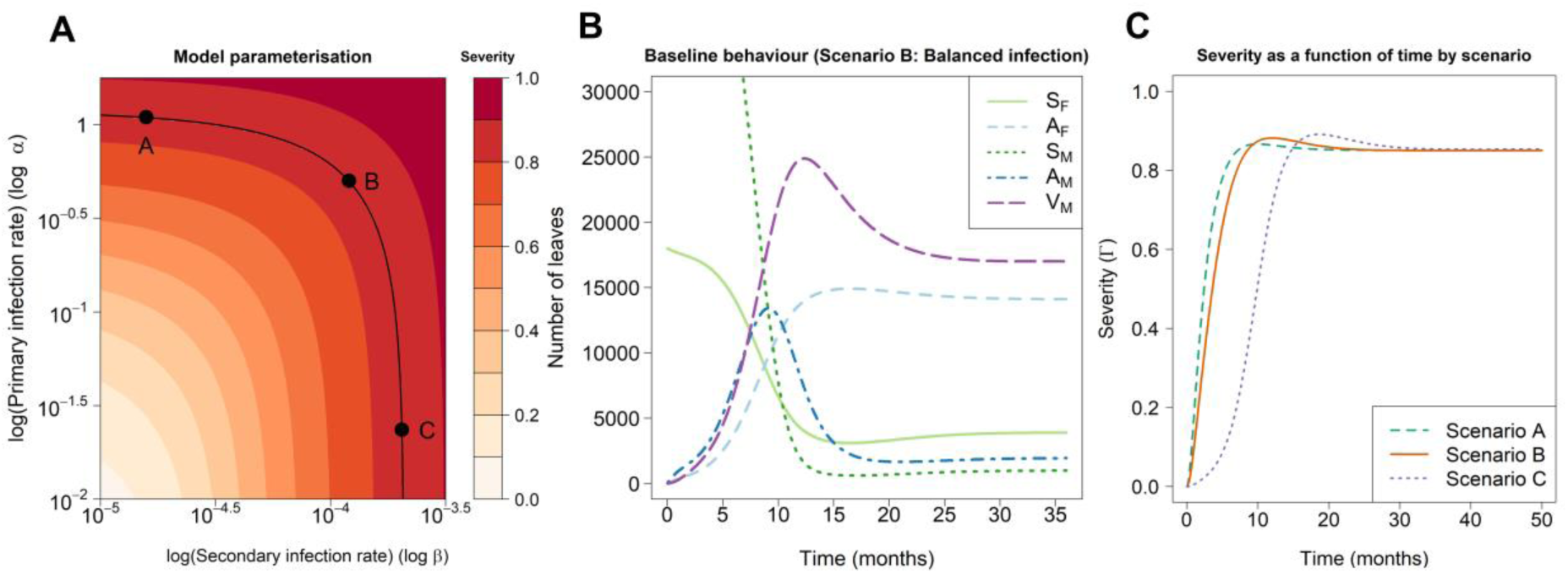
Parameterisation of the single-grower model. (A) Terminal disease severity as a function of the primary (ε) and secondary (β) infection rates (each axis is on a log10 scale). Pairs of values leading to severity 0.85 – which roughly matches our disease spread data (cf. Fig. 4) – are linked with the black curve. Three parameter scenarios are identified and marked with black dots: Scenario A (high primary infection but low secondary infection), Scenario B (balanced primary and secondary infection) and Scenario C (low primary infection but high secondary infection). (B) Baseline behaviour of the model in Scenario B, showing the numbers of leaves in each compartment on the average tree within a grove for three years after first introduction of the disease into a grove of mature trees. (C) Severity as a function of time under each parameterisation scenario.

From this set of possible parameters, we identify three representative pairs of values corresponding to the following three illustrative scenarios.

#### Scenario A (“Primary-dominated”)

High Primary, Low Secondary, with ε ≈ 1.1 month^-1^ and β ≈ 1.6 × 10^−5^ month^-1^, in which most infection is caused by sources external to the grove of interest.

#### Scenario B (“Balanced infection”)

Balanced Primary and Secondary, with ε ≈ 5.0 × 10^−1^ month^-1^ and β ≈ 1.2 × 10^−4^ month^-1^, in which the epidemic is driven roughly equally from sources within the grove *vs.* external to the grove.

#### Scenario C (“Secondary-dominated”)

Low Primary, High Secondary, with ε ≈ 2.3 × 10^−2^ month^-1^ and β ≈ 2.1 × 10^−4^ month^-1^, with most infection coming from within the grove of interest.

Disease reaches equilibrium following first introduction in a mature grove rapidly under all three scenarios (shown in Fig. 9B for the balanced infection scenario). At equilibrium the rate of production of new susceptible flush is equal to the sum of the outflows of flush due to leaf maturation and flush leaf infection, with the rate of flush maturation equal to sum of the rates of mature leaf infection and mature leaf natural death (*cf.* Fig 2; the sum of flows into and out of each compartment are equal, for each compartment). While there are differences in the initial response of severity to time between scenarios (Fig. 10C), reflecting a transition from “monomolecular-like” (primary-dominated scenario) to “logistic-like” (secondary-dominated scenario) disease progress curves (Madden *et al*. 2007), differences are relatively slight. Any distinction between the scenarios is certainly not captured in the data we have taken here, since the disease was already very well-established in the groves we studied. This reiterates the idea that further experimental work would be required to conclusively disentangle the relative rates of the two infection pathways.

**Figure 10.**
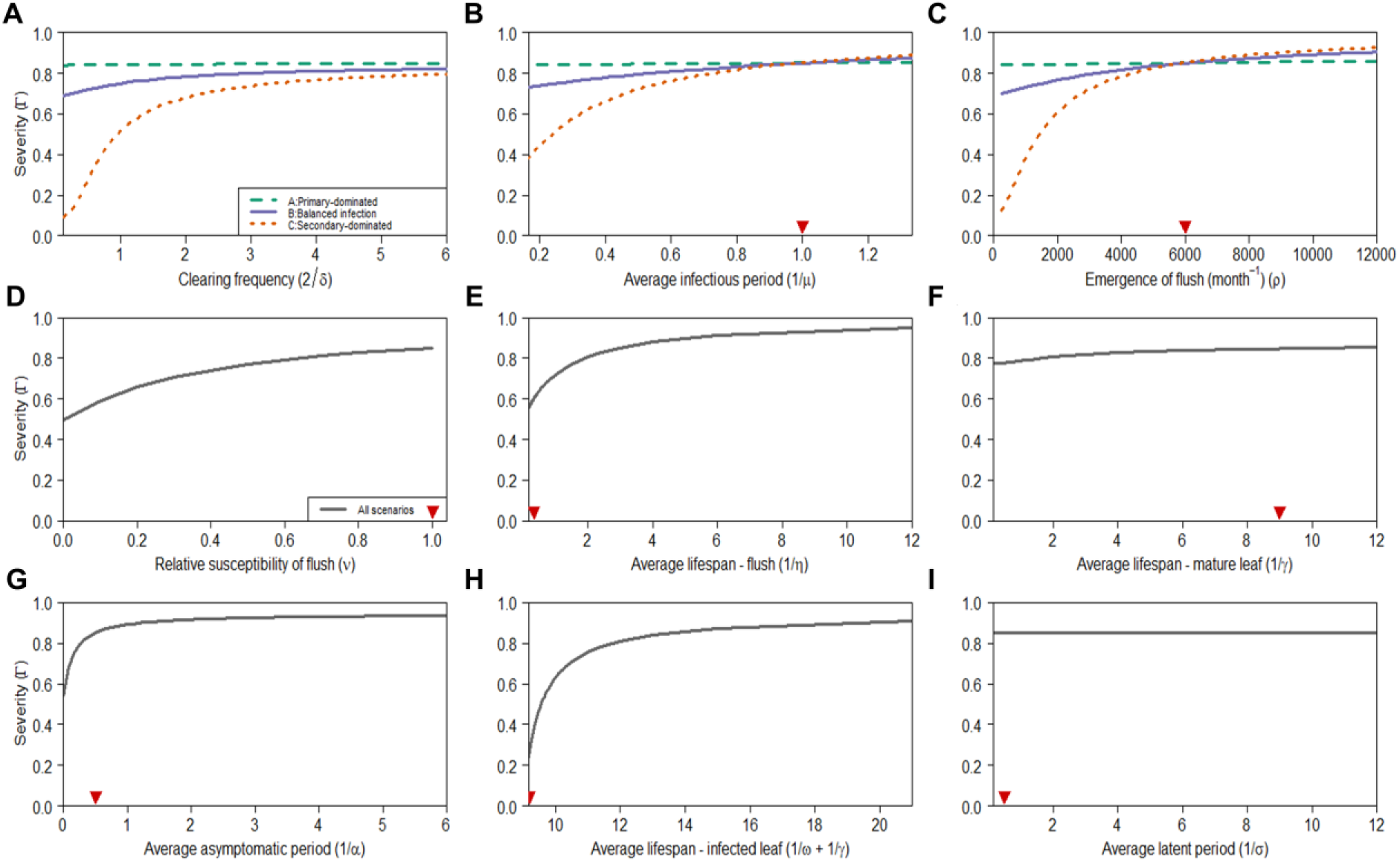
Sensitivity Analysis: impact of parameter values on equilibrium severity. (A) Interval between successive clearances of leaf litter. (B) Average lifespan of diseased leaf (without control). (C) Birth rate of flush. (D) Relative susceptibility of flush to mature leaves. (E) Average age of maturity (“lifespan”) of flush. (F) Average lifespan of mature leaves in a disease-free system. (G) Average asymptomatic period. (H) Average infectious period of fallen leaves. (I) Average latent period. Red arrows on the x-axes show default parameterisation. All periods are in months. The parameter scenario only affects the responses panels A-C, in which cases the scenarios are distinguished by different types and colours of line; in panels D-I the responses in all three parameter scenarios are co-incident.

### Modelling disease control

Although not captured in our data, the relative balance of primary and secondary infection can have strong implications upon which types of control can be successful (Bassanezi *et al.*, 2013; Bergamin Filho *et al.*, 2016). To explore this further, we consider removal of fallen leaf litter. Following the logic concerning roguing given in Cunniffe *et al*. (2014), if the leaf litter is removed every Δ months, then the average fallen leaf remains on the ground for Δ/2 months, and so – assuming that all leaves are removed on each round of litter removal – the appropriate rate of removal in our model is δ = 2/Δ.

### Sensitivity analysis of the single-grower model

We performed a sensitivity analysis to examine how changes in parameter values affected terminal disease severity in the single-grower model across each of the primary-infection dominated, the balanced and the secondary-infection dominated scenarios. The response of the equilibrium severity did not vary between scenarios for the majority of the epidemiological parameters we considered (Figs. 10D–I), since the effects of these parameters do not interact with the rates of infection. Most patterns were intuitive. For example, the severity decreases as the relative susceptibility of flush (ν) decreases (Fig. 10D). We note, however, that it is unsurprising that terminal severity did not vary with the average latent period for leaf litter (1/σ; Fig. 10I), since the default parameterisation for the rate of removal of litter δ is 0. As it is never cleared, the litter will always become infectious and thus contribute to terminal severity, irrespective of the latent period on the ground.

Responses to three parameters, however, did vary across infection rate scenarios. An increased birth rate of flush (ρ; Fig. 10C) has its greatest impact on the secondary-dominated scenario. Similarly, short infectious periods (1/μ; Fig. 10B) caused substantial reductions in terminal severity in the secondary-dominated scenario, though infectious periods >2 months resulted in little change from the baseline parameterisation (1 month). The changes in terminal severity for the primary-dominated and balanced infection scenarios were less severe. In the single-grower version of the model, there is no dependence of rates of primary infection on the amount of infection present within the grove, so changes in infectious periods have little impact on the amount of infection coming from these sources.

Considering the response of the equilibrium severity to different frequencies of litter removal (Fig. 10A) reveals significant differences in the predicted effectiveness of control based on the relative balance of primary and secondary infection. In particular, under Scenario A, the high rate of inoculum ingress from outside the grove means that purely localised control based on local removal of litter has very little effect on the disease severity, no matter how frequently the litter is removed. Even very high rates of local litter removal under Scenario B – in which both primary and secondary infection have a role in driving the epidemic dynamics – also do not greatly affect the level of disease (e.g. weekly leaf removal leads to a reduction in terminal severity from 85% to around 70%). However unsurprisingly in Scenario C – in which local inoculum almost entirely drives the epidemic – local removal of leaf litter can be very effective, even if only practised by the single grower in question.

### Area-wide disease management

While the results given in Fig. 10A provide a useful baseline, disease control by removal of leaf litter is more likely to be more successful if adopted by an entire community of growers, since this would reduce sources of inoculum for all groves. We demonstrate this using our model of area-wide control (Fig. 11), assuming that the rate of primary infection, ε, depends on the proportion of growers (π) practicing control (see Methods). The performance of disease management within the set of groves doing control depends strongly on the parameter scenario (Figs. 11A,D,G), with the results shown in Fig. 10A providing an upper bound on the severity when no one else clears leaf litter (i.e. when π = 0). Similarly, the results shown in Fig. 10A provide a lower bound on severity for growers not doing control (Figs. 11B, E, H). In all parameter scenarios, removal of litter every month or so is required for eradication of disease if there is 100% compliance. However, the performance averaged over all growers when there is an intermediate level of compliance (Figs. 11C, F, I) depends on the parameter scenario. For example, monthly removal of litter by 75% of growers leads to average severities over all groves of 0.56, 0.50 and 0.27 in our primary-dominated, balanced infection and secondary-dominated parameter scenarios, respectively.

**Figure 11.**
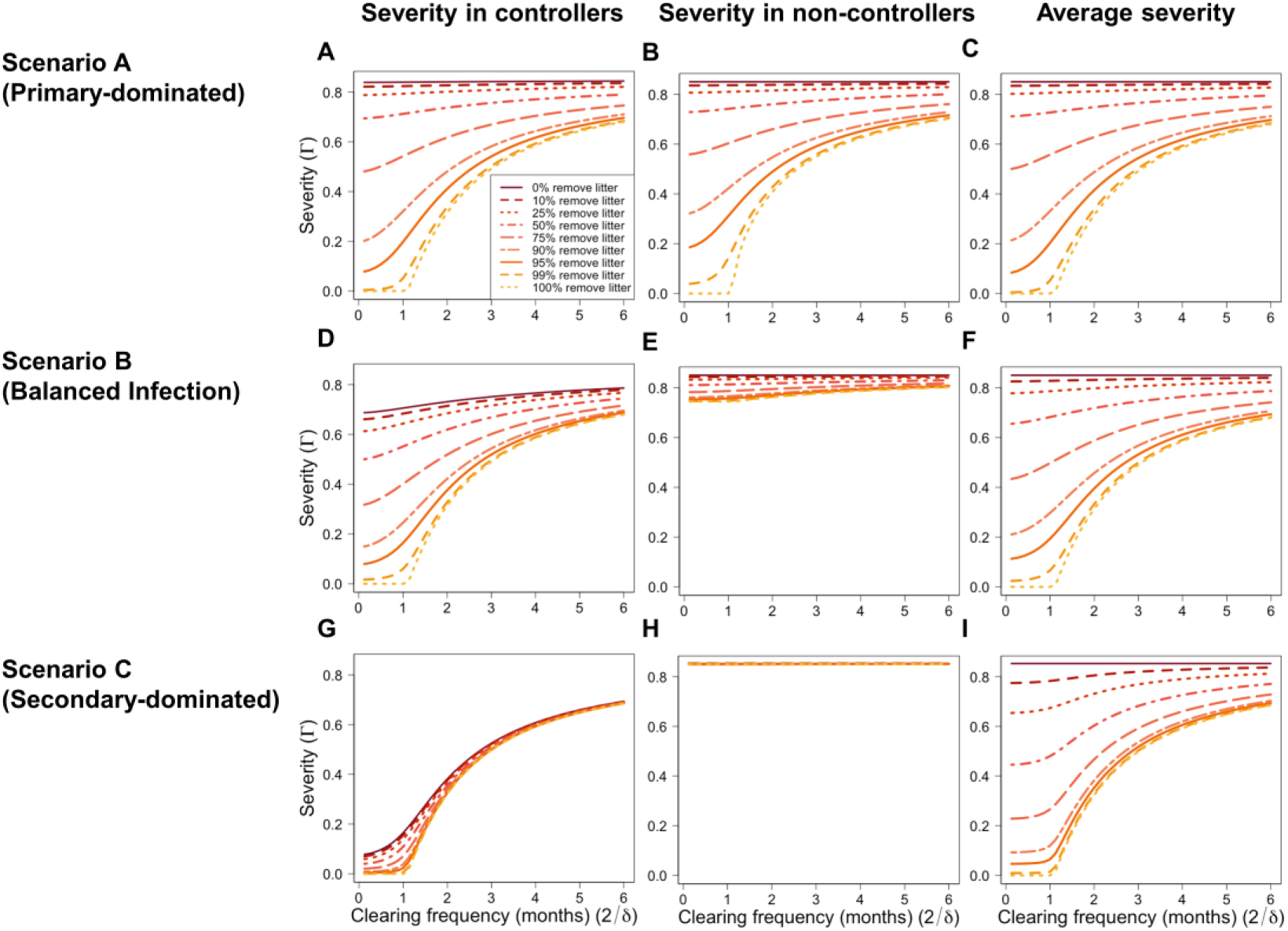
Area-wide disease management. The impact of a proportion □ of grower controlling disease by local removal of leaf litter (at average frequency 2/δ) on disease severity in groves of controllers (Γ_*C*_; panels A, D and G), non-controllers (Γ _*NC*_; panels B, E and H) and on average over all groves (πΓ_*C*_ + (1− π)Γ_*NC*_; panels C, F and I). The top row (panels A-C) shows the primary-dominated parameter scenario (Scenario A); the middle row (panels D-F) shows the balanced infection parameter scenario (Scenario B), and the bottom row (panels G-I) shows the secondary-dominated parameter scenario (Scenario C).

### Frequency of litter removal for eradication

Area-wide eradication requires all growers to participate (i.e. π = 1). Our numerical work suggests it also requires growers to remove leaf litter at least once a month or so, and that this minimum removal frequency does not depend on the parameter scenario. This can be explained using the basic reproduction number (*R*_*0*_). In particular, in the area-wide disease spread model, when π = 1 and so when all growers control disease, the basic reproduction number is given by

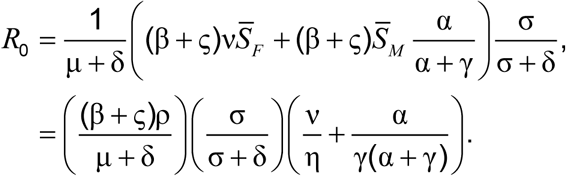

The expression can be understood by considering the number of infections caused by a single infected leaf introduced into the disease-free system, within which the numbers of susceptible flush and mature leaves are 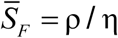 and 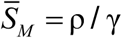, respectively. The initially infected leaf remains infectious for 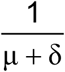 units of time. It infects flush leaves at net rate 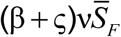 and mature leaves at net rate 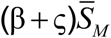. Any (asymptomatic) infected flush leaf definitely becomes a symptomatic mature leaf, whereas an asymptomatic infected mature leaf becomes symptomatic before being abcised with probability 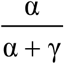 (since it has to develop symptoms and so move to the *V*_*M*_ compartment before falling to the ground). In both cases mature leaves detach to become uninfectious fallen leaf litter, and become infectious before decaying with probability 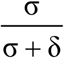. This completes a single infection cycle: combining the various rates and probabilities leads to the expression for *R*_0_ given above.

The expression for *R*_0_ is a decreasing function of the litter clearance rate δ, which – for all parameterisations of the model (Table 1) – is below 1 if δ > 1.81 per month. Given the relationship δ = 2/Δ, this correponds to litter removal every Δ = 2/1.81 ∼ 1.1 months, i.e. the threshold obtained in all three parameter scenarios for the area-wide model (Figs. 11C,F,I). The particular parameter scenario is not important in setting this threshold because the value of (β + ς) is fixed at precisely the same value, i.e. that which leads to a terminal severity of 0.85 in the single-grove model in the absence of disease management (the numeric value is (β + ς) ≈ 2.1×10^−4^ month^-1^).

## Discussion

CGS symptoms were found in every single grove and plant, irrespective of sub-region, altitude, grove age or variety. This pattern was consistent over time for each of the ten sampled groves in Cruz das Almas, and so, at this spatial level (i.e. the region), CGS can be considered to be highly regularly dispersed. Our data therefore confirm the conclusion of the preliminary report of Silva *et al*. (2009) that CGS in Reconcavo is currently endemic in the sense that it is regularly found and very common in that particular area.

The level of *Z. citri* infection is more related to conditions for epiphytic growth than to the number of ascospores (Mondal & Timmer, 2003b). We did not find evidence that CGS intensity in Recôncavo Baiano is higher on the lower canopy as was previously found in Florida (Mondal *et al.*, 2003), perhaps since *psl* was so high for all heights. This can perhaps be linked to the negative logarithmic pattern for the number of trapped *Z. citri* ascospores at different heights reported in Florida; even at 7.5m some spores were captured (Mondal *et al.*, 2003). Nevertheless, considering that the closest CGS inoculum source is ground level litter (Mondal & Timmer, 2006b), lack of variability over heights throughout regions indicates that conditions are conducive for *Z. citri* dispersion/infection and that CGS is very widespread.

The ubiquity of CGS was confirmed for almost all sampling units during the monthly evaluations in Cruz das Almas groves. The increase in *psl* was somewhat expected, as most reports show such pattern for Cuba, Costa Rica and Florida (Garcia *et al.*, 1980; Hidalgo *et al.*, 1997; Mondal & Timmer, 2003a). However, as those studies were primarily concerned with the development of CGS symptoms on new shoots, they reported a strong seasonality, and had incidences ranging from as low as 30% to as high as 100%. Here, as we focused on the presence of CGS on mature rather than on new leaves, we could evaluate what we consider the real incidence for the Recôncavo region, reflecting the epidemiology of the pathogen rather than the demography of its host.

Our distributed lag analysis supported no link between disease severity and abiotic environmental drivers. In Florida it was shown that the less intense epiphytic growth occurred in dry months, with growth speed increasing during the summer (Mondal & Timmer, 2003a). In Cuba, an increase in CGS incidence and severity during Summer was reported (rains and high relative air humidity) (Garcia *et al.*, 1980). However, in Recôncavo da Bahia climatic conditions appear to provide no obstacle to CGS, with no extreme temperatures, as well as long periods of high relative air humidity and only short periods without rain. Such conditions are ideal for the *Z. citri*-citrus interaction.

Since no variability was found in the fourth and third levels of the spatial hierarchy (i.e. among municipalities or groves), they were not analysed. When such analyses were feasible (i.e. among plant quadrants and plants), the index of dispersion indicated a random pattern for single evaluations and at both spatial levels. However, the general spatial pattern was regularity (log(*A*)<0 and *b*<1) as shown by the binary power law. This favours the hypothesis that allo-infections (i.e. from inoculum coming from outside the grove) might be as important as auto-infections. This strengthens the case for area-wide management.

Spore trapping experiments could help to assess the allo-infection hypothesis. Another way of confirming the relative balance of auto- and allo-infection might be to collect data at lower disease severity levels, which could be used to determine the rates of primary and secondary infection ε and β in our single-grower mathematical model (cf. Fig. 9A). Targetted experimentation, e.g. performing litter removal in some experimental plots while allowing disease to progress unhindered in others, would be another way to begin to disentangle the balance of the routes of infection. However, very large plots might be required to obtain a detectable effect. Such data would also allow us to verify the time-dynamics of our model, and in particular to verify that the time-progression in the number of leaves in each compartment over time is plausible (*cf.* Fig 10). However, we do not have such data in hand.

Given this uncertainty concerning the balance of the infection pathways, we restricted ourselves to a relatively simple mathematical model of the system, replicating only the important features of the *Z. citri* infection cycle, with all parameters taking constant values. Nevertheless, by systematising available knowledge on CGS epidemiology, our model allows us to understand potential effects of manangement. In particular, we have shown that cultural control – of the type that could be done by resource-poor growers in Northeastern Brazil – might potentially be successful. However, assuming primary infection is indeed at least somewhat implicated in CGS epidemiology, area-wide management would almost certainly be required (Bassanezi *et al.*, 2013; Bergamin Filho *et al.*, 2016) (Fig. 11). Conclusions would be similar for any localised control affecting only secondary within-grove infection, e.g. accelerating the decay of inoculum by treating litter with urea (Mondal & Timmer, 2003a).

In our single-grower model we assumed a constant rate of primary infection. This assumption is often made in models (e.g. Cunniffe *et al*., 2015), including in studies fitting models to data (see e.g. Parry *et al*., 2014). We later extended the model to include the behavior of other growers and the effects of area-wide control on any individual’s local epidemic. We then assumed that the rate of primary infection depends on the average level of infection area-wide. Even without full participation, our model showed concerted efforts across growers lead to large reductions in disease severity for growers who manage disease.

Rates of primary infection will also be affected by regional environmental conditions, as well as the spatial structure of the landscape. Here we ignored spatial structure and assumed homogeneity amongst growers of the same type. However, relaxing this condition would be an interesting extension to the model. The relative balance of primary infection to secondary infection would also vary depending on the size of the grove of interest (Hilker *et al.*, 2017), and its location within the landscape. Finally, in the area-wide model we assumed all primary infection was caused by inoculum produced within the set of citrus groves tracked. This allowed disease to be eradicated when all growers control disease. In practice some inoculum might come via very long-distance dispersal from outside the area of interest, or be produced on other sources such as non-cultivated citrus within the region. Our results are therefore an upper bound upon the levels of disease control that could be achieved. If other sources of inoculum do in fact exist, divergence from this optimal performance would be most significant for the scenario in which primary infection dominated (i.e. Scenario A).

*Z. citri* is already highly dispersed in Recôncavo of Bahia, with maximum CGS prevalence and incidence in plants and their quadrants, irrespective of the location of the groves or plant’s age. This fact, associated with the high incidence in leaves seems a natural consequence of the way in which CGS has been viewed in that region; because it has never been considered important, it was never controlled. Hence, the incidence increased, but as this process has been cyclic and apparently quite slow, the disease is still not seen as a threat (Rodrigues, 2018). The situation resembles that previously in Florida, where prior to 1940 CGS was not considered a serious problem (Mondal & Timmer, 2006a). Only when growers finally noted the disease was causing defoliation, did they begin to attempt control.

In light of our results, CGS can be regarded as endemic in Recôncavo da Bahia. Moreover, in Bahia state the disease is also found in other regions (unpublished). In contrast with the situation in São Paulo, the most important citrus producing region in Brazil, many diseases are not present in Recôncavo. For instance, this area is currently free from HLB (*Candidatus* Liberibacter spp.), leprosis (CiLV), citrus canker (*Xanthomonas citri* subsp. c*itri*) and sweet orange scab (*Elsinöe australis*) (Laranjeira *et al.*, 2005). In this context, at least for the region we are concerned with, CGS cannot be regarded as a minor citrus disease. Nevertheless, growers face a temporal continuity of both host (perennial crop with multiple flushings per year) and pathogen (putative abundant inoculum due to favourable weather and multiple inoculum sources) coupled to a host spatial continuity represented by more than 10,000 ha of citrus in the region (IBGE, 2017). Hence, due to its endemic status CGS control in Recôncavo will be a challenging task. Some growers try to invest in technology, but overall low input level farming represents the citrus industry. Citrus groves are established without long term planning and simple small-holder farming takes place. Growers often have a low educational level and low concern about general horticultural practices, soil management or plant protection. However, there is no evidence of different CGS intensity or perception according to groves’ technological level (Rodrigues, 2018).

CGS control in the Recôncavo region will probably demand a set of practices at a frequency and cost which is incompatible with the current technological level and economic position of local citrus growers. Moreover, due to the abundance of inoculum, favourable weather and long-distance wind-dispersion of ascospores, localised control attempts in small groves may have a low probability of success. Therefore, if growers decide to tackle this disease, it is advisable that their efforts should be coordinated on a regional basis. Additional data would allow us to confirm this preliminary conclusion, as well as to continue to model the system. Collecting more disease spread data and using these data to parameterize more detailed mathematical models will form the basis of our future work on this pathosystem.

## Acknowledgements

The authors would like to thank Mr Decio de Oliveira Almeida for technical support, and Dr Matthew Castle, Dr Adam Hall and Mr Elliott Bussell for useful discussions. FFL and ACFS are fellows of Conselho Nacional de Desenvolvimento Científico e Tecnológico (CNPq). The authors thank the anonymous reviewers and the editors for comments which were instrumental in greatly strengthening the final paper. Conceived and designed experiments: SXBS, FFL. Provided funds and materials: SXBS, FFL, HPSF. Performed the experiments: SXBS. Analyzed the data: SXBS, FFL. Designed the model: FFL, NJC. Analysed the model: NJC, REMW, FFL. Wrote the paper: FFL, NJC, REMW, SXBS. Contributed to drafting the manuscript: FFL, NJC, REMW, SXBS, ACFS. Part of this paper is from a MSc dissertation presented by SXBS to the Universidade Federal do Recôncavo da Bahia.

## SUPPLEMENTARY TEXT 1

We examined data on the number (out of 20) of symptomatic leaves at each of 3 heights on 30 plants in each of 3 groves.

The data were therefore as follows

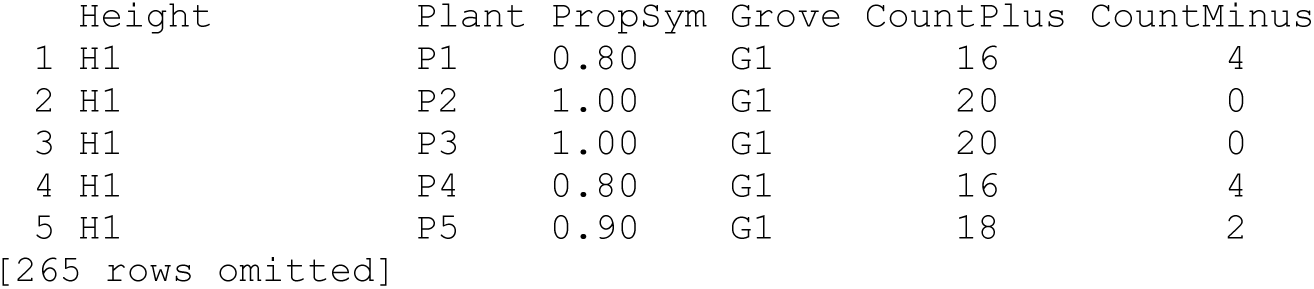

(there were 270 such measurements, for 3 heights on 30 plants in 3 groves [i.e. 90 plants in total]).

The raw data are plotted below. It suggests that there might be an increase in severity with height in Grove 1, but that any effect is more marked in Groves 2 and 3.

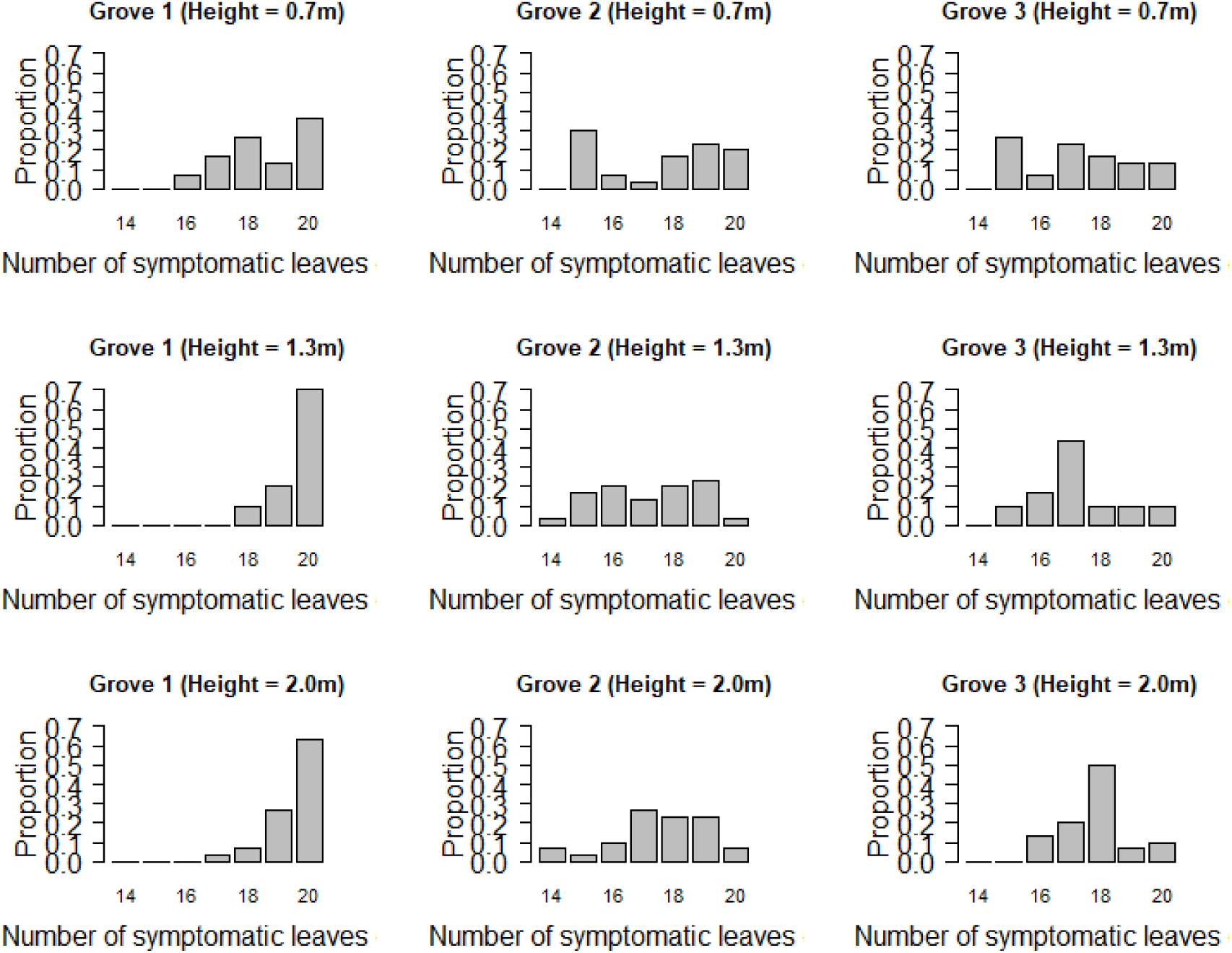

The mixed effect model we used to analyse these data uses a random effect for each plant to focus on the differences between disease severities on the same plant at different heights.

These are plotted overleaf; note that the histograms in rows two and three of the following figure plot out the differences on a plant-by-plant basis between the counts of infected leaves at H2 = 1.3m (row two) and H3 = 2.0m (row three) relative to H1 = 0.7m.

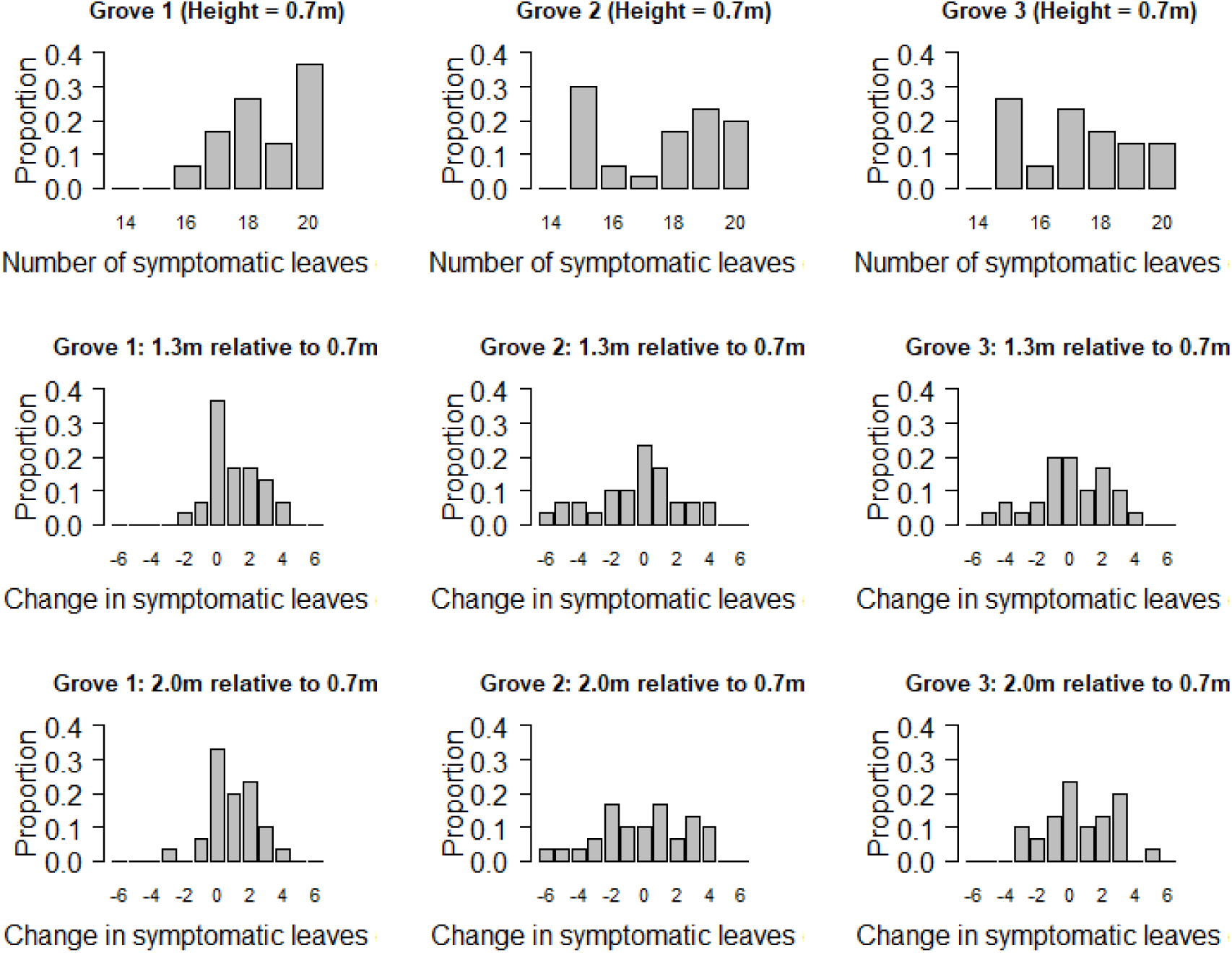

The effect of plant height on disease severity appeared equivocal between the three groves we tested; there appears to be an increase in Grove 1, but no clear pattern in Groves 2 and 3. This preliminary visual conclusion was supported by our statistical modelling. We fitted a mixed effect logistic regression and found statistically support for an interaction between the two fixed effects `Height` and `Grove` (in a model with a random intercept for each `Plant`).

~~~
glmer(cbind(CountPlus,CountMinus) ∼ Height + Grove + Height:Grove + (1 | Plant),
family = binomial(link = “logit”), data = CGS_data)
~~~

The plot below shows the odds at different heights for different groves in this best-fitting model.

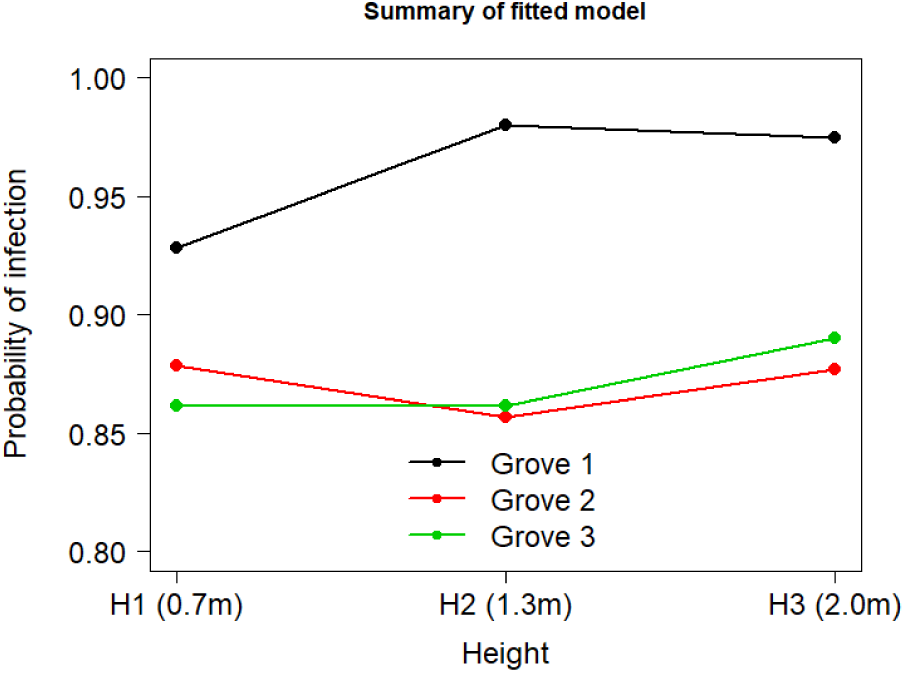

## Notes

https://github.com/nikcunniffe/Citrus-Greasy-Spot

